# Environmental controls of dark CO2 fixation in wetland microbiomes

**DOI:** 10.1101/2024.01.18.576062

**Authors:** Luise Grüterich, Jason Nicholas Woodhouse, Peter Mueller, Amos Tiemann, Hans-Jo-achim Ruscheweyh, Shinichi Sunagawa, Hans-Peter Grossart, Wolfgang R. Streit

## Abstract

Rising atmospheric concentration of CO_2_ is a major concern to society due to its global warming potential. In soils, CO_2_ fixing microorganisms are preventing a part of the CO_2_ from entering the atmosphere. Yet, the pathways behind dark CO_2_ fixation are rarely studied *in situ*. Here we examined the environmental controls on the abundance and expression of key genes involved in microbial CO_2_ fixation in estuarine wetlands. A combined multi-omics approach incorporating metabarcoding, deep metagenomic and metatranscriptomic analyses confirmed that wetland microbiota harbor all six known CO_2_ fixation pathways and that these pathways are transcribed at high frequencies along several environmental gradients, albeit at different levels depending on the environmental niche. Notably, the transcription of the key genes for the reductive tricarboxylic acid cycle (rTCA) and the Calvin cycle were favored by low salinity and O_2_ rich niches high in organic matter, while the transcription of the key genes for the Wood-Ljungdahl pathway (WLP) and dicarboxylate/4-hydroxybutyrate cycle (DC/4-HB cycle) were favored by low O_2_ niches poor in organic matter. Taxonomic assignment of transcripts implied that dark CO_2_ fixation was mainly linked to few bacterial phyla, namely, Desulfobacterota, Gemmatimonadota, Methylomirabilota, Nitrospirota and Pseudomonadota.

## INTRODUCTION

Rising atmospheric concentrations of the greenhouse gases (GHGs) CO_2_ and methane are the primary drivers of climate change and hence a major concern to society. Therefore, knowledge on fundamental processes linked to biological CO_2_ producing and -fixing approaches is crucial (1, 2). Within this framework, current research focuses on GHG fixing microbial metabolic pathways that could be relevant to the climate system because of their ability to counteract the increasing atmospheric GHG concentrations.

Light-driven photosynthesis and the associated CO_2_ fixation pathways (i.e. Calvin cycle) are well understood and play an obvious role in the climate system (3, 4).

In addition to light-dependent CO_2_ fixation, also non-photosynthetic GHG fixation pathways can play a major role in the climate system. One prominent example of non-photosynthetic microbial GHG fixation is methanotrophy. Methanotrophs (aerobic and anaerobic methane oxidizers) are responsible for preventing up to 90% (the number varies depending on the ecosystem) of the methane produced in soils or aquatic systems from entering the atmosphere (5–7). Extensive, targeted research has refined our quantitative and mechanistic understanding of methanotrophy and its role in the global climate system (6, 8, 9). By contrast, non-photosynthetic microbial CO_2_ fixation (dark CO_2_ fixation), beyond hydrogenotrophic methanogenesis, has received much less attention by the scientific community. However, it could be similarly important as methanotrophy for balancing the climate system. Studies on dark CO_2_ fixation were almost exclusively conducted in single-species pure cultures (e.g. (10, 11)). Thus, their role in the global climate systems remains largely unknown. Within this setting in addition to the well-known Calvin cycle, light independent CO_2_ fixing pathways have more and more come to the attention of researchers (10, 12). Among the domains of life, at least six different autotrophic CO_2_ fixation pathways exist (Calvin cycle, rTCA cycle, WLP, 3-hydroxypropionate bicycle (3-HP bicycle), 3-hydroxypropionate/4-hydroxybutyrate cycle (3-HP/4-HB cycle) and DC/4-HB cycle) (13–15) (TABLE 1). It was recently reported that high levels of CO_2_ drive the

**Table 1:**
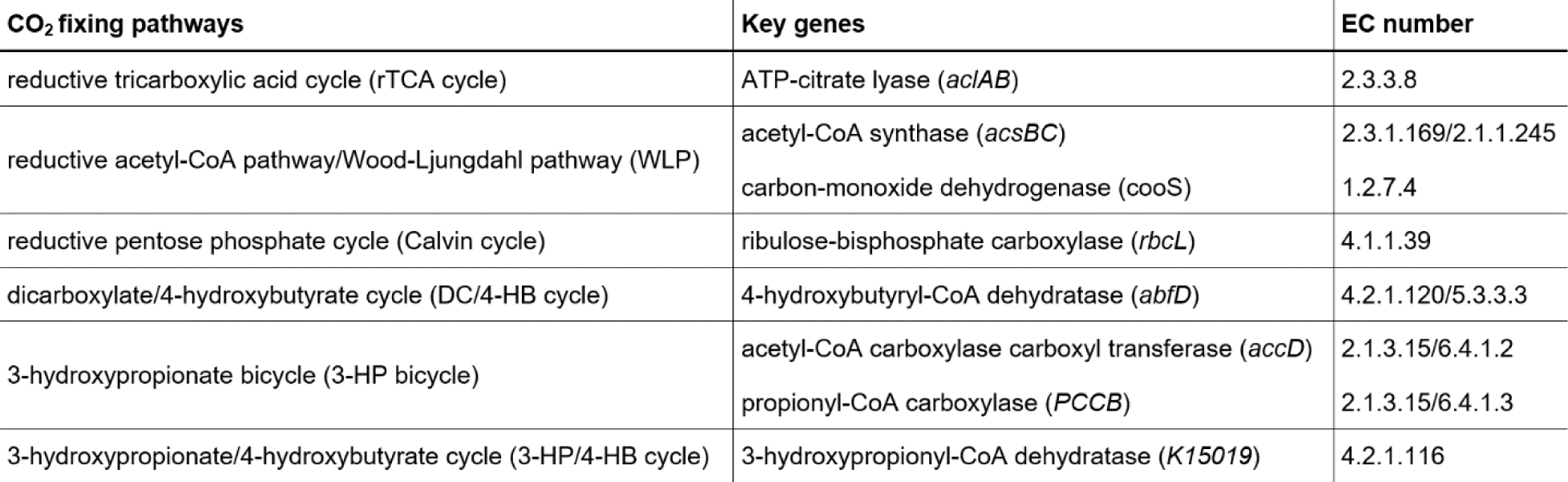
List of CO_2_ fixing pathways and affiliated key genes.

TCA cycle backwards towards autotrophy (12). Due to the increase in CO_2_ concentration in the atmosphere mainly caused by anthropogenic fossil fuel burning, it is now important to elucidate the environmental factors that determine occurrence and magnitude of the respective dark CO_2_ fixation pathways *in situ*.

The present study assessed the potential of the different dark CO_2_ fixing pathways along steep environmental gradients. The estuarine wetland landscape is ideally suited to understand environmental constraints on CO_2_ fixing pathways for at least two reasons. First, estuarine wetlands connect marine, limnic, and terrestrial environments, and are therefore characterized by steep environmental gradients in O_2_, organic matter availability, and salinity. Second, estuarine wetland soils possess a tremendous carbon turnover rate and constitute at the same time among the most effective carbon sinks of the biosphere (16, 17). Our study was conducted along three wetland sites, i.e. tidal marsh ecosystems, along the salinity gradient of the Elbe estuary, NW Germany. The Elbe estuary with a coverage of 500 km^2^ is one of the largest estuaries in Europe with various environmental gradients. For this study, we considered gradients in salinity, flooding frequency and soil depth at three sampling sites ranging from marine to freshwater marshes (FIGURE 1).

**Figure 1:**
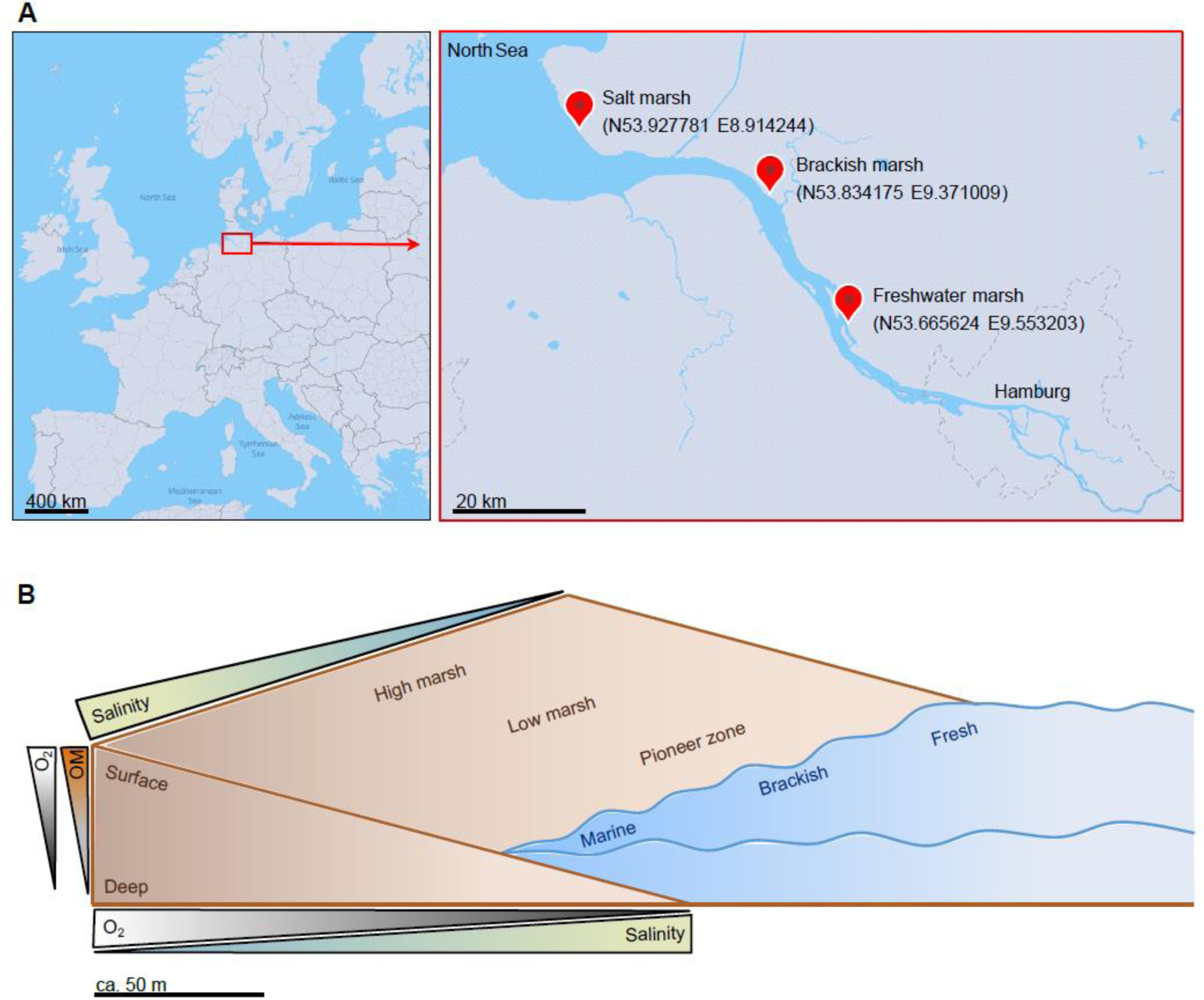
Location of the Elbe estuary and sampling sites along the salinity gradient of the Elbe estuary from the freshwater marsh via the brackish marsh to the salt marsh (A). B shows the environmental gradients given by the sampling locations along the river course from fresh via brackish to marine sites, along the flooding frequency gradient from high marshes via low marshes to pioneer zones and the different depths we sampled (surface layer (0 cm) to the deep layer (50 cm)). At each site, in each zone and depth, samples were taken with a peat sampler in September at low tide. Maps in A were created with MapTiler.

We hypothesized (I) that dark CO_2_ fixation has an ecological niche in anaerobic environments and increases with decreasing O_2_ availability, because many of the CO_2_ fixing pathways have been described in anaerobic microorganisms (13). Anaerobic environments thus may represent an ecological niche, where dark CO_2_ fixation might be competitive in relation to heterotrophic microbial metabolism, but (II) also profit from heterotrophic respiration of CO_2_ that we expect to increase with increasing salinity, which is associated with a greater availability of alternative electron acceptors such as sulfate, facilitating microbial respiration of CO_2_ under anoxic conditions (18, 19).

While many studies have examined the abundance and expression of key genes for CO_2_ fixation pathways in specific ecosystems or under specific environmental conditions, this study investigates expression across large environmental gradients, allowing for identification of niches of these pathways. Analyzing key genes targets crucial, pathway-specific genes, which are unique to the respective microbial pathways. We provide strong evidence that salinity, flooding frequency, and soil depth do affect the expression level of dark CO_2_ fixation. This observation, based on metagenome-, metatranscriptome- and metagenome assembled genome (MAG) analysis, is supported by metabolite- and phylogenetic analyses.

## RESULTS

In order to constrain the ecological niches of dark CO_2_ fixing pathways *in situ*, the soil microbiome was assessed along salinity, flooding and soil-depth gradients. We expected several environmental parameters to covary along these gradients, namely, organic matter availability in relation to soil depth, O_2_ availability in relation to soil depth and flooding frequency, and the availability of alternative electron acceptors (e.g. sulfate) – and thus respiratory capacity under anoxic conditions – in relation to salinity and flooding frequency.

We assessed the extent to which wetland microbiota harbor the potential for dark CO_2_ fixation, and how gene abundance and expression vary along environmental gradients in salinity, flooding frequency and soil depth (FIGURE 1). In total, our sampling design includes 69 amplicon data sets (FIGURE 2) as well as 18 metagenome and the corresponding metatranscriptome data sets (FIGURE 4) with associated organic matter and metabolite data (FIGURE S1). All underlying samples were collected in September. Thus, all resulting data represent a snapshot at this time of year.

**Figure 2:**
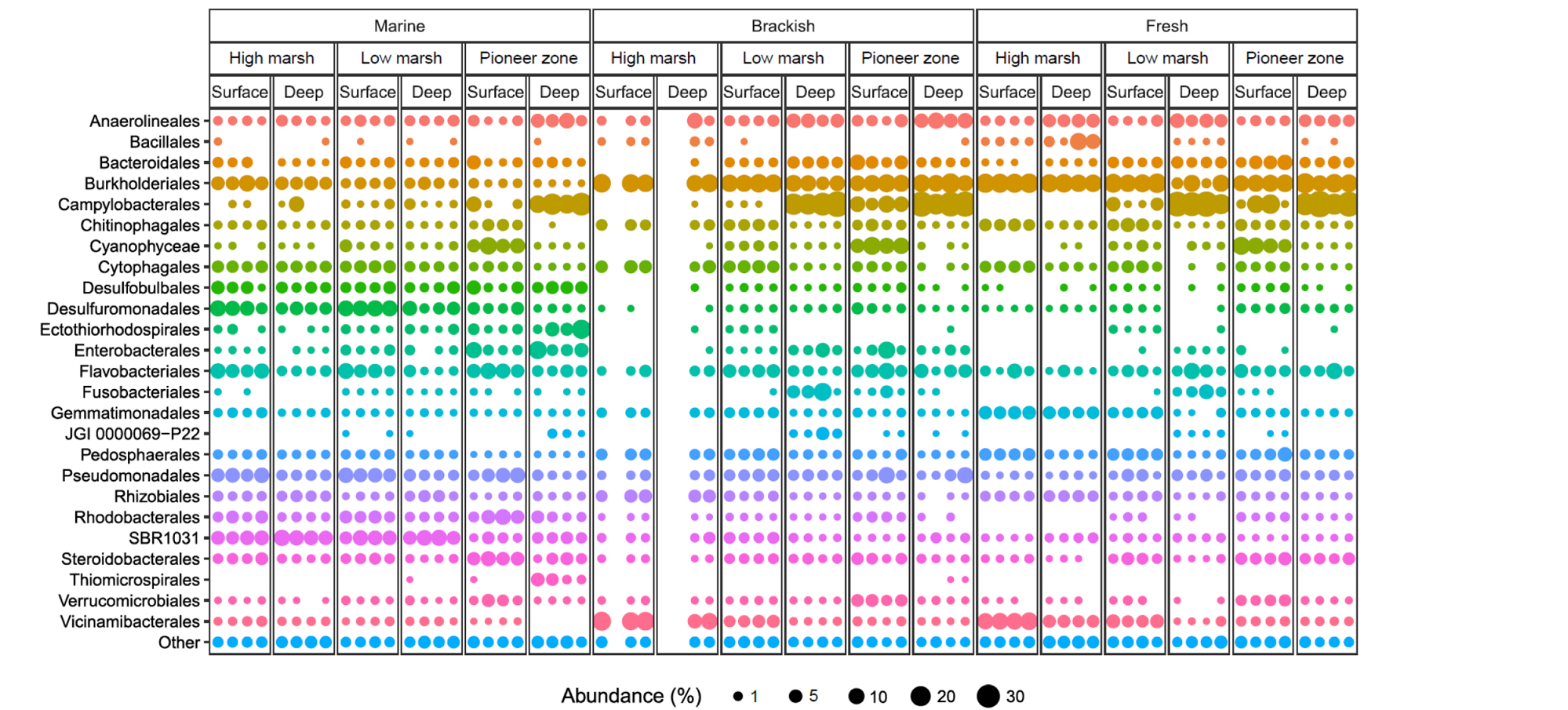
Taxonomic composition of bacterial communities at the order level for the 18 sampling locations with four biological replicates resulting in a total of 69 analyzed samples (three replicates could not be included due to insufficient sequence quality). Samples originate from the three sites: Salt marsh (marine), brackish marsh (brackish) and freshwater marsh (fresh). Each from the three zones: High marsh, low marsh and pioneer zone. Each from the surface layer at 0 cm (surface) and the deep layer at 50 cm depth (deep). Depicted are the 25 orders with the highest average relative abundances. All other orders were combined into the “Other” group.

### Phylogenetic indications for microbial CO_2_ fixation

Phylogenetic analysis was performed to obtain further insight into the metabolic capabilities of wetland microbiota (20, 21). Samples were taken as soil cores at 18 locations and DNA and RNA were extracted as specified in Material and Methods. The 16S rRNA amplicon sequencing results of all 69 samples indicated that the top four most abundant orders were Campylobacterales with an average of 10.2, Burkholderiales with an average of 9.7%, Flavobacteriales with an average of 3.9% and Anaerolineales with an average of 3.3% (FIGURE 2). Campylobacterales, Burkholderiales and Anaerolineales harbor species capable of CO_2_ fixation (22–24). Comparing the abundance of the 25 most abundant orders, shifts in community composition occur along the different environmental gradients (FIGURE 2).

16S rRNA provides the advantage of observing multiple functional group shifts simultaneously. However, connecting phylogenetic groups to function is challenging due to microbial plasticity and functional redundancy. Limited knowledge exists about the impact of microbial composition on community-level physiological profiles (25).

### Metagenomics show high potential for microbial CO_2_ fixation in estuarine wetlands

The above-made analyses strongly suggest a high metabolic diversity of the marsh soil microbiota. While many of the identified orders harbor species capable of CO_2_ fixation (22–24), we wanted to assess the genetic potential for dark CO_2_ fixation along environmental gradients (FIGURE 1). For this we sequenced 18 metagenomes. For each sample a minimum of 54 million and an average of 99 million reads were obtained with an average N50 value of 1,173 (TABLE S1). A schematic overview of the bioinformatics workflow is provided in figure 3.

**Figure 3:**
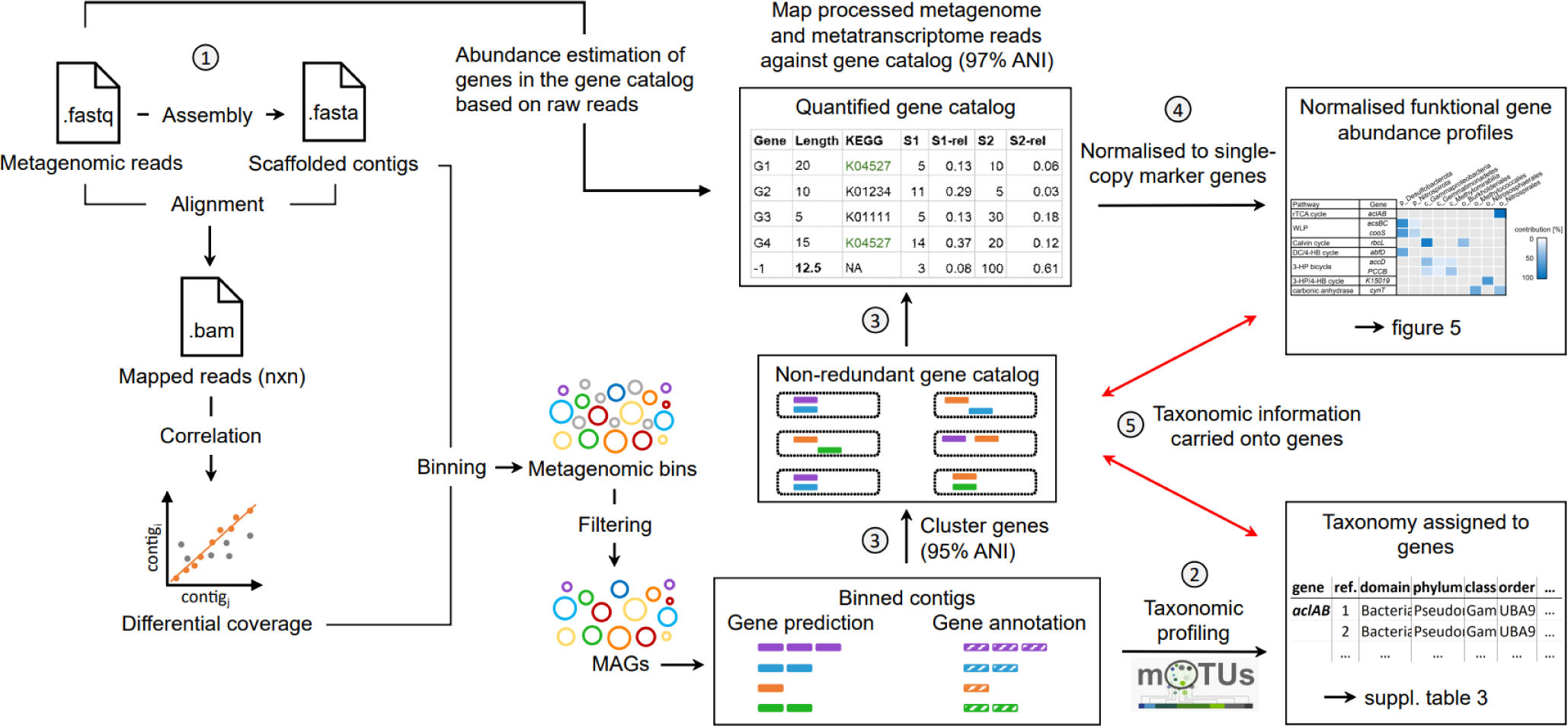
Schematic overview of reads to gene catalog pipeline. 1) Processed reads from each sample were assembled independently, and all samples were used to guide per sample binning 2) MAGs were placed into species level mOTUs using single copy marker genes, and assigned taxonomy using GTDB-TK, used for accurate taxonomic profiling. 4) Processed reads were mapped against the 3) Gene prediction and annotation were performed on all prokaryotic MAGs, and a non-redundant gene catalogue was built using a clustering threshold of 95% ANI. 4) Processed metagenome- and metatranscriptome reads were mapped against the gene catalogue and normalized based on gene length (within sample normalization) and single-copy marker gene abundance (between sample normalization). 5) Information linking taxonomy from MAGs and gene/transcript abundance was maintained at all times to allow for taxonomic interrogation of functional gene abundance.

On the basis of the obtained metagenome data, we analyzed the relative abundance of key genes for the different metabolic pathways involved in microbial CO_2_ fixation. For this, we determined the mean gene copy number per assembled genome of nine well-known genes involved in the six known microbial CO_2_ fixation pathways. Although we are interested in dark CO_2_ fixation, for comparison, we also included the gene encoding for RuBisCO (*rbcL*), which is the key gene in the Calvin cycle, and the gene for the carbonic anhydrase (*cynT*), which mediates the CO_2_ supply for metabolic pathways by solubilizing CO_2_ and forming HCO_3_ ^−^ (FIGURE 4A, TABLE 1).

**Figure 4:**
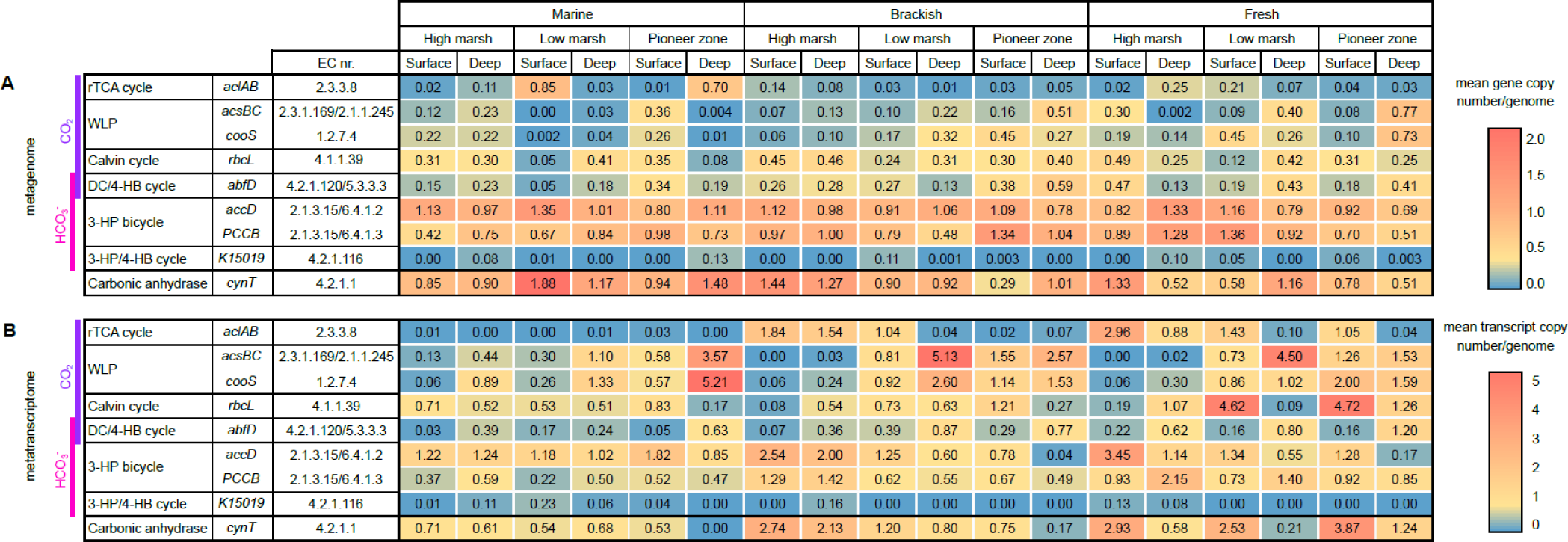
Metagenome and metatranscriptome analysis of genes involved in CO_2_ fixation. Abundance of the key genes (copy number per genome) (A) ATP-citrate lyase (*aclAB*), acetyl-CoA synthase (*acsBC*), carbon-monoxide dehydrogenase (*cooS*), ribulose-bisphosphate carboxylase (*rbcL*), 4-hydroxybutyryl-CoA dehydratase (*abfD*), acetyl-CoA carboxylase carboxyl transferase (*accD*), propionyl-CoA carboxylase (*PCCB*), 3-hydroxypropionyl-CoA dehydratase (*K15019*) and carbonic anhydrase (*cynT*) and their respective transcripts normalized to single copy marker gene (B). The purple bar indicates the pathways that use CO_2_ directly as carbon source and the pink bar those that use HCO_3_^−^ as carbon source. The 18 depicted samples originate from the three sites: Salt marsh (Marine), brackish marsh (Brackish) and freshwater marsh (Fresh). Each from the three zones: High marsh, low marsh and pioneer zone. Each from the surface layer at 0 cm (Surface) and the deep layer at 50 cm depth (Deep).

We were able to detect the presence of all nine genes involved in CO_2_ fixation, although with varying copy numbers along the environmental gradients (FIGURE 1B). The copy number of the nine genes investigated ranged from a mean gene copy number/genome of 0 to 1.88. Compared to the mean gene copy number/genome of the Calvin cycle key gene (*rbcL*), which ranged from 0.08 to 0.49, the relative gene abundances of the key genes assigned to CO_2_ fixation in the dark are even higher for some pathways (FIGURE 4A).

The gene linked to the 3-HP/4-HB cycle (*K15019*) had the lowest abundance of the nine investigated key genes with a mean gene copy number of 0 to 0.23 per genome. The most frequently detected genes were linked to the 3-HP bicycle (*accD* and *PCCB*), which had copy numbers ranging from 0.42 to 1.36. Similarly high copy numbers were detected for carbonic anhydrase (*cynT*) (FIGURE 4).

On the genomic level, only three significant (p < 0.05) changes along the three environmental gradients (i.e., salinity, flooding, soil depth) could be observed. Gene abundance of the key gene for the DC/4-HB cycle (*abfD*) decreased with increasing salinity, gene abundances of *accD* decreased with increasing flooding frequency and the WLP key gene (*acsBC*) increased with increasing flooding frequency (FIGURE 4A; TABLE 2A).

**Table 2:**
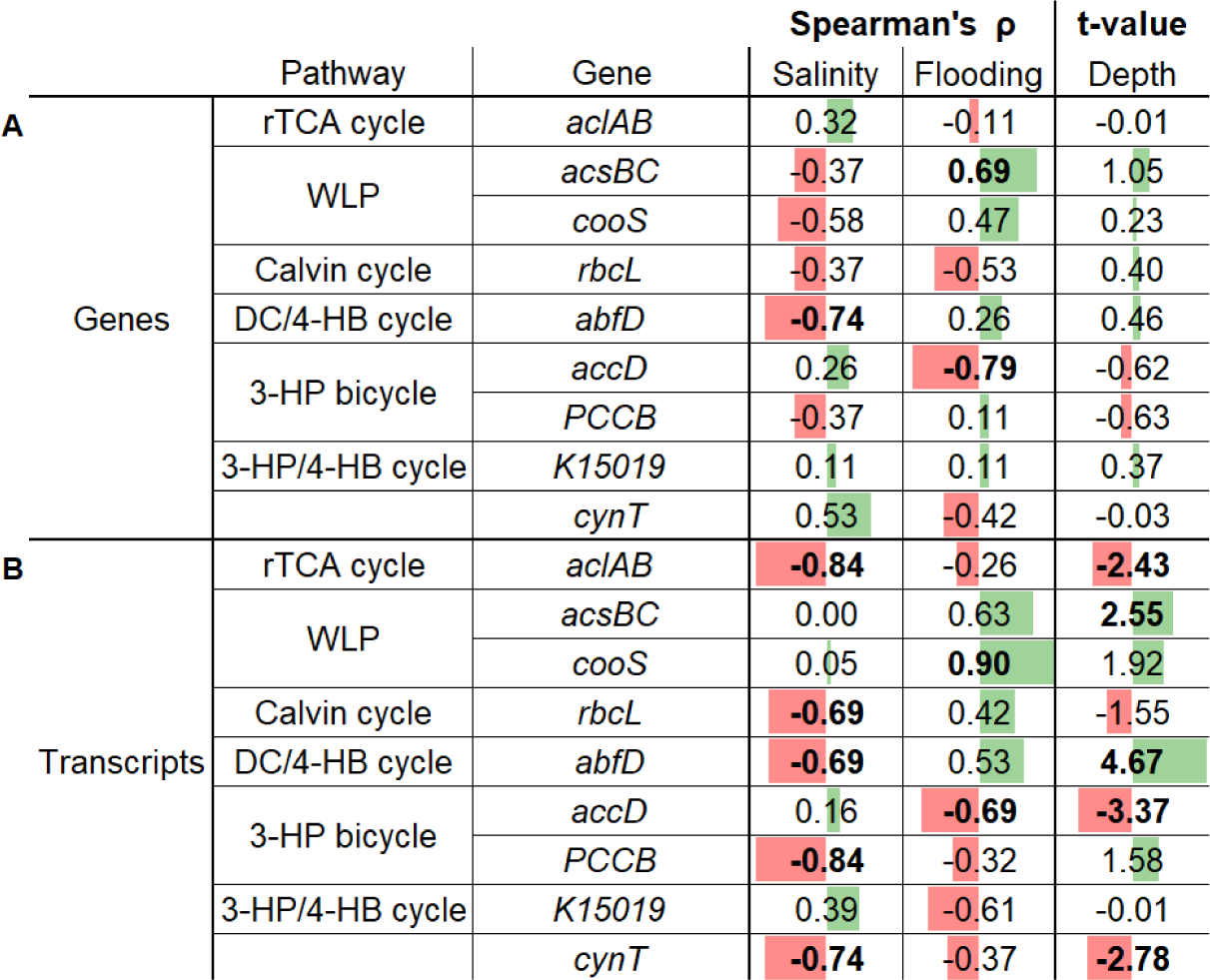
Statistical analysis of the genes and transcripts of the nine key genes related to CO_2_ fixation. Significant changes are indicated by bold numbers (p < 0.05). For changes along the salinity gradient and the flooding frequency gradient, we performed a Spearman’s rank correlation. For changes along the depth gradient, we performed a paired t-test. For the column “Salinity” we have increasing abundance with increasing salinity (green) and decreasing abundance with increasing salinity (red). For the column “Flooding” we have increasing abundance with increasing flooding frequency (green) and decreasing abundance with increasing flooding frequency (red). For the column “Depth” we have increasing abundance with increasing soil depth (green) and decreasing abundance with increasing soil depth (red).

### Metatranscriptomics reveal high expression levels of key genes related to microbial CO_2_ fixation

Intrigued by the observation that genes for all CO_2_ fixing pathways were present along the investigated environmental gradients (FIGURE 1, 4A), we asked to what extend these genes were transcribed. Therefore, we analyzed the transcription level of the key genes involved in CO_2_ fixation (TABLE 1, FIGURE 4B).

RNA from a total of 18 soil samples was extracted as described in the Material and Methods section, used for mRNA sequencing and mapped to the key genes (TABLE 1, FIGURE 4B). For each sample a minimum of 50 million and an average of 80 million reads were obtained with an average N50 value of 750 (TABLE S1). We detected transcripts of all nine CO_2_ fixation affiliated key genes. The number of mapped transcripts varied strongly along the environmental gradients (FIGURE 1, 4B). Transcripts ranged from a mean transcript copy number/genome of 0 to 5.21 (FIGURE 4B).

On the transcriptomic level we found several significant changes (p < 0.05) along the environmental gradients (FIGURE 4B, TABLE 2B). This finding clearly contrasts against the metagenomic results. Along the salinity gradient, we observed a significant decrease (p < 0.05) of transcripts with increasing salinity for the key genes of the rTCA cycle (*aclAB*), Calvin cycle (*rbcL*), DC/4-HB cycle (*abfD*) and 3-HP bicycle (*PCCB*) and for the carbonic anhydrase (*cynT*). Along the flooding frequency gradient, we observed a significant decrease (p < 0.05) of the WLP key gene (*cooS*) with decreasing flooding frequency. The opposite was true for the key gene of the 3-HP bicycle, the transcript abundance of which increased with decreasing flooding frequency. Along the soil depth gradient, a significant decrease (p < 0.05) of transcripts with increasing depth was observed for the key genes of the rTCA cycle (*aclAB*) and the 3-HP bicycle (*accD*) and the carbonic anhydrase (*cynT*). Significant increases (p < 0.05) of transcripts with soil depth were observed for the key genes of the WLP (*acsBC*) and DC/4-HB cycle (*abfD*) (FIGURE 4B, TABLE 2B). Among all investigated genes, the key gene of the 3-HP/4-HB cycle (*K15019*) showed the weakest expression along all gradients (FIGURE 4B).

For some genes the high transcript abundance was confined to specific environmental niches. However, gene expression level did not correlate with the number of observed gene copies. For instance, the highest number of WLP transcripts was observed in the deep layer of the salt marsh pioneer zone, the deep layer of the brackish low marsh and the deep layer of the freshwater low marsh, which harbored rather low numbers of respective gene copies (FIGURE 4).

The fact that the results of metagenomics and metatranscriptomics clearly differed, indicated that the combined approach was essential for understanding the environmental constraints on the extent of dark CO_2_ fixation.

Based on our MAG analysis, we were able to assign function to taxonomy (FIGURE 5, TABLE S3, S4). Average abundances were calculated based on the samples, where the transcript abundance assigned to the respective microbial group was higher than 0. Because of excluding the zero values, abundances cannot be just added up.

**Figure 5:**
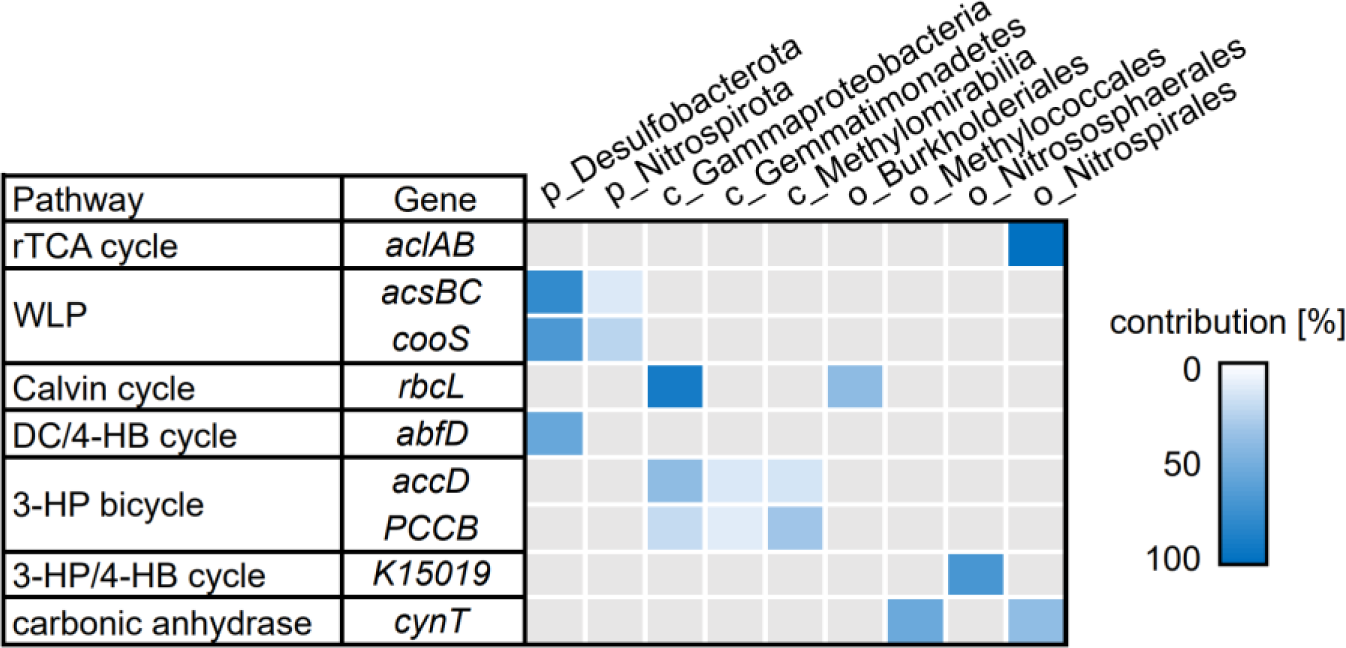
Most abundant microbial groups assigned to the nine genes associated with dark CO_2_ fixation based on MAGs. Blue color intensities indicate the average relative abundance of the microbial groups (phylum-(p), class-(c) or order level (o)) of all 18 samples, where the transcript abundance assigned to the respective microbial group was higher than 0. Because of excluding the zero values, abundances should not be added up. Underlying values and per-site abundances are shown in TABLE S4.

An intriguing observation was that transcripts for all key genes were predominantly associated with a limited number of microbial groups. The rTCA cycle was primarily driven to an average of 99.8% by the order Nitrospirales. Main drivers behind the WLP were the phyla Desulfobacterota (72.7%) and Nitrospirota (16.5%). The transcripts of RuBisCO were predominantly assigned to the class Gammaproteobacteria (91.1%), with the order Burkholderiales forming the majority of Gammaproteobacteriales with an average abundance of 38.3% across all environmental gradients. Transcription of the DC/4-HB cycle was mainly driven by the phylum Desulfobacterota, with an average abundance of 55.2%. The primary driving classes of the 3-HP bicycle were Gammaproteobacteria (28.3%), Gemmatimonadetes (10.4%), and Methylomirabilia (22.1%). The archaeal order Nitrososphaerales played a major role in the few transcripts of the 3-HP/4-HB cycle, with an average abundance of 68.9%. The orders responsible for transcripts of carbonic anhydrase (*cynT*) were Nitrospirales (37.3%) and Methylococcales (53.4%).

## DISCUSSION

Phylogenetic and metagenomic analyses identified both heterotrophic and chemo- or photo-autotrophic microorganisms. The main orders observed were Campylobacterales, Burkholderiales, Flavobacteriales and Anaerolineales of which Campylobacterales, Burkholderiales and Anaerolineales are known to harbor species capable of CO_2_ fixation (22–24). Furthermore, it is known that the genetic potential for dark CO_2_ fixation is spread over a broad taxonomic range (26). In general, for all sites, the diversity of 85 different phyla and 185 different classes indicated a highly diverse metabolic potential of the microbiota (TABLE S2) considering there is a total of 89 bacterial phyla known to the Silva database (27). Metagenome sequence analyses showed that the key genes of all known CO_2_ fixation pathways are present in the marsh soils of the Elbe estuary, suggesting a major potential for microbial CO_2_ fixation. Our metatranscriptomic analyses supported the expression of these pathways, although we observed a disconnect between gene copy numbers and expression levels. Indeed, a high number of transcripts can be observed, although gene abundance is very low. We, therefore, refer to changes in CO_2_ fixing pathways along the environmental gradients based on the expression, rather than gene copies, of key genes.

Our sampling design covered steep gradients in organic matter availability in relation to soil depth, O_2_ availability in relation to soil depth and flooding frequency, and availability of alternative electron acceptors (e.g. sulfate) in relation to salinity and flooding frequency.

Our hypotheses that (I) increasing salinity and (II) decreasing O_2_ availability increase dark CO_2_ fixation could only partly be confirmed. Most pathways show a significant response (p < 0.05) in terms of gene expression to soil depth and salinity while the gene expression for only two genes showed a significant response (p < 0.05) to changes in the flooding frequency gradient (TABLE 2). The expression of the rTCA cycle, Calvin cycle and the carbonic anhydrase was favored by low salinity and the soil surface environments (FIGURE 4B, TABLE 2B).

Even if the rTCA cycle is restricted to anaerobic or microaerophilic organisms (28), we showed that it is not restricted to strictly anaerobic environments that we would expect in the deep soil layers. Our findings indicate a preference of the rTCA cycle to high marsh environments, which are only occasionally anoxic due to a low flooding frequency (flooding restricted to storm tides) (FIGURE 1, TABLE 2B; (29, 30)) and characterized by greater organic matter availability supporting higher rates of heterotrophic microbial activity and stimulation of CO_2_ fixation, which was correlated with respiration in previous studies (FIGURE S1; (26, 30–32)). Due to the reversibility of the TCA to the rTCA cycle (10), microorganisms can easily switch to the rTCA cycle if needed, especially in generally aerobic soils, when forced to switch from aerobic to anaerobic metabolism during occasional events of anoxia. Surprisingly, at the two locations with the highest gene abundance, no gene expression was observed. The order behind the highest level of transcription of the rTCA key gene (*aclAB*) was Nitrospirales, contributing 98-100% of the *aclAB* transcripts varying from site to site. *Nitrospira* sp. Palsa-1315 was the dominating genus behind *aclAB* transcripts (FIGURE 5, TABLE S3, S4). It belongs to the clade B Nitrospira (33) and so far, no pure culture of clade B Nitrospira exists, therefore physiological responses to environmental controls need to be further established (34). Our transcriptomic results show, that the genus *Nitrospira* sp. Palsa-1315 only occurs in brackish to freshwater environments.

The gene abundance and transcription of the Calvin cycle’s key gene RuBisCO is not strictly dependent on the availability of light although highest transcript abundance was recorded in two surface layer samples. For other wetland ecosystems, Burkholderiales was found to be one of the dominant photosynthetic order (23). In our samples, for all sites, that showed transcription of RuBisCO, an average of 38% of the transcripts, also originate from photosynthetic bacteria of the order Burkholderiales (FIGURE 5, TABLE S3, S4).

Not all CO_2_ fixing pathways use CO_2_ in its atmospheric form as a carbon source. In porewaters, depending on pH, inorganic carbon occurs in various species, namely, CO_2_, carbonate ions (CO ^2−^) and bicarbonate ions (HCO_3_ ^−^) (35). The potential of the enzymatic reaction from CO to HCO ^−^ is highly favored by the presence of the carbonic anhydrase (*cynT*). Other studies suggest, that the activity of *cynT* is correlated with the environments with rich plant diversity (36). Our data suggest that transcripts of *cynT* gene homologs are favored by the O_2_- and organic matter rich soil-surface and low-salt environments (FIGURE 4B, TABLE 2B).

The expression of the WLP and DC/4-HB cycle was favored by deep soil environments and high flooding frequency (FIGURE 4B, TABLE 2B). Regarding acetogenesis represented by the WLP, the *cooS* gene reflects the same ecological niche as the *acsBC* gene. This can be expected because, together, they form the key enzyme complex of the WLP (37). We observed that expression of WLP key genes (*acsBC* and *cooS*) appears to be linked to the deep soil layers and frequently flooded zones (FIGURE 4B). We therefore conclude that low O_2_ availability is the driving environmental factor favoring WLP activity because both deep soil layers and regularly flooded zones are low-O_2_ environments. Recent transcriptome studies in brackish water microbial mats showed that Desulfobacterota are capable of expressing the WLP and subsequent acetate and pyruvate metabolism (38). This can be supported by our taxonomic assignment of the *acsBC* and *cooS* transcripts, which show that Desulfobacterota play an important role in the expression of WLP key genes (FIGURE 5, TABLE S3, S4). We also observed accumulation of pyruvate in the environmental niches, that harbor high numbers of *acsBC* and *cooS* transcripts indicating that the WLP as source of acetate could lead to pyruvate production by the pyruvate formate-lyase (39) which show a very high transcript abundance in our samples ranging from 0 to 31 mean transcripts/genome (FIGURE S1).

The DC/4-HB cycle was, like the WLP, also favored in the deep soil layers and frequently flooded zones (FIGURE 4B, TABLE 2B). The gene expression of its key gene (*abfD*) seems therefore also be favored by environments of low O_2_ availability, presumably, because carboxylation is performed by the O_2_-sensitive pyruvate synthase and phosphoenolpyruvate carboxylase (40).

## CONCLUSION AND PERSPECTIVE

Our study demonstrates that wetland microbiota are drivers for dark CO_2_ fixation. Wetlands cover only 1% of the Earth’s surface, but store about 20% of the total ecosystem organic carbon stock (41). Their global contribution to dark CO_2_ fixation might therefore be of high relevance to the climate system. This study decrypts the influence of salinity, flooding and soil-depth gradients as environmental factors that control the expression of dark CO_2_ fixation in wetland ecosystems.

This study shows, a) that only five microbial groups (the phyla Desulfobacterota and Nitrospirota and the classes Gammaproteobacteria, Gemmatimonadetes and Methylomirabilia) are responsible for the majority of dark CO_2_ fixation assigned transcription. b) Both community composition and the abundance and expression of functional genes showed high variability in relation to environmental gradients within the estuarine wetland landscape. c) Our findings suggest, that the expression of the rTCA cycle, the Calvin cycle and the carbonic anhydrase, was driven by O_2_-, organic matter- and salinity. d) For the expression of the WLP and DC/4-HB cycle, we argue that O_2_ is the strongest driver behind niche occupancy. e) Variations in the expression of specific functional genes across environmental gradients were more prominent than variations in microbial groups or genes related to dark CO_2_ fixation, suggesting that functional diversity plays a crucial role by allowing microorganisms to adapt to their specific environment, leading to niche formation.

Concerning our metatranscriptomic analyses it is important to note that although transcriptomes are a good tool to demonstrate the actual expression of a gene, it must be considered that this is always only a snapshot at the time of sampling. Therefore, in further studies, a temporal component, at a small scale to cover the tidal cycle or at a large scale to cover seasonal effects, should be considered in addition to the spatial level. Our study aimed at determining the drivers of dark CO_2_ fixation pathways along environmental gradients, and, for the majority of pathways, we succeeded at identifying important environmental controls. However, our study cannot determine the relative importance of dark CO_2_ fixation pathways in terms of counterbalancing ecosystem GHG production. There is an increasing number of studies indicating negative CO_2_ fluxes in the dark in many ecosystem types such as saline/alkaline soils, dry inland waters and dry river sediments (42–44). Future research will need to incorporate biogeochemical tools to understand the relative importance of microbial dark CO_2_ fixation in the climate system.

## MATERIAL AND METHODS

### Soil sample collection

Soil samples were collected from three wetland sites along the salinity gradient of the Elbe estuary (September 2021 and September 2022): a salt marsh (N53.927781° E8.914244°), a brackish marsh (N53.834175° E9.371009°) and a freshwater marsh (N53.665624° E9.553203°). Samples were collected with a peat sampler (Eijkelkamp, Giesbeek, Netherlands) to a depth of 50 cm below the soil surface in each of three elevation zones at each site, the high marsh, low marsh and pioneer zone. Aliquots of each soil sample were stored at −80 °C until use for DNA/RNA extraction and metabolite analysis.

### DNA and RNA extraction from soil

DNA was extracted from aliquots of all collected soil samples, which were frozen at −80°C until analysis. Upon thawing on ice, 0.5 g of soil were used for DNA extraction using the NucleoSpin Soil Kit (Macherey-Nagel, Düren, Germany) following the manufacturer’s protocol. Subsequently, the isolated DNA was analyzed at a wavelength of 280 nm using a Nanodrop spectrophotometer (NanoDrop 2000, Thermo Scientific, Waltham, USA). 2 g of soil were used for RNA extraction using the RNeasy PowerSoil Kit (Qiagen, Venlo, Netherlands) following the manufacturer’s protocol. RNA concentrations were quantified using the Qubit 2.0 Fluorometer and the RNA High Sensitivity Assay Kit (RNA HS, Thermo Fisher, Berlin, Germany).

### Determination of organic matter via Loss of Ignition method

The weight change related to high-temperature oxidation of organic matter was used to determine the organic matter content. About 30-40 g of soil was placed in porcelain crucibles and dried overnight at 60 °C in a drying. Before ignition the samples were dried for 2 h at 105 °C in a drying oven to get rid of remaining water. The dry weight was measured. The soil samples were ignited for 2 h at 500 °C in a muffle furnace, cooled in a desiccator and weighed again. The g organic matter (OM)/g dry weight (DW) was calculated as follows:

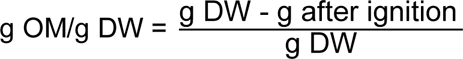

### Metabolite analyses via colorimetric assays

Acetate, pyruvate and isocitrate concentration was determined by colorimetric assay kits that were used according to the respective manufacturer’s protocol (Sigma; MAK086, MAK071, MAK061). 1 g of soil was mixed with 1 ml of VE H_2_O, followed by ultrasonication and subsequent centrifugation at 11000 g for 1 min. The supernatant was used as sample for metabolite determination.

### 16S rRNA gene analyses

Metabarcoding sequencing of 16S rRNA variable regions V3-4 was performed at the Competence Centre for Genomic Analysis, Kiel, Germany using the Illumina Nextera XT Index Kit and primers 341F (5’-CCTACGGGNGGCWGCAG-3’ and 785R (5’-GACTACHVGGG-TATCTAATCC-3’) and MiSeq Reagent Kit v3. Adaptor trimming of demultiplexed paired-end reads was performed using Cutadapt (v.4.4) (45). Read filtering and ASV inference was performed using the DADA2 pipeline (v.1.8), with the following specifications (maxEE=2, maxN=0, truncQ=2, truncLen=260,210). Taxonomic assignment of merged corrected reads was made using the SILVA (v.138.1) database. Downstream analyses were performed using the phyloseq (v.1.41.1) and ggplot2 (3.4.2) R packages.

### Meta-omics sequencing and data processing

Size selection, library preparation and sequencing of metagenomes were performed at the Ramaciotti Centre for Genomics, Sydney, Australia, using the Ilumina DNA Prep kit and an Illumina NovaSeq 6000 S4 flow cell. Ribosomal RNA depletion, library preparation and sequencing of metatranscriptomes was performed at the Competence Centre for Genomic Analysis, Kiel, Germany using the Illumina Stranded Total RNA Prep with Ribo-Zero Plus Microbiome kit and Illumina NovaSeq 6000 S4 Reagent kit.

Our data analysis, which has been previously demonstrated (46, 47) set a new benchmark for metagenomic and metatranscriptomic analyses. As MAGs are only partial representations of a population and pathways associated with individual MAGs may not always follow the same dynamics as the MAGs themselves, this approach emphasizes quantification at the level of a non-redundant gene catalog that is derived from MAGs and maintains taxa-function information. Further because complex metagenomes can be influenced by the heterogenous occurrence of eukaryotes, viruses and other genetic elements, rather than normalizing to the size of the library (i.e. FPKM, RPKM metrics), this approach normalizes the abundance of individual genes against prokaryotic single-copy marker genes allowing us to estimate the copies per prokaryotic cell.

Metagenomic (n=18) and metatranscriptomic (n=18) sequencing datasets were processed as previously described in (46). Briefly, BBMap (v.38.71) was used to quality control sequencing reads from all samples by removing adapters from the reads, removing reads that mapped to quality control sequences (PhiX genome) and discarding low-quality reads (trimq=14, maq=20, maxns=1, and minlength=45). Quality-controlled reads were merged using bbmerge.sh with a minimum overlap of 16 bases, resulting in merged, unmerged paired, and single reads. The reads from metagenomic samples were assembled into scaffolded contigs (hereafter scaffolds) using the SPAdes assembler (v3.15.2) (48) in metagenomic mode. Scaffolds were length-filtered (≥ 1000 bp) and quality-controlled reads from each metagenomic sample were mapped against the scaffolds of each sample. Mapping was performed using BWA (v0.7.17-r1188; -a) (49). Alignments were filtered to be at least 45 bp in length, with an identity of ≥ 97% and a coverage of ≥ 80% of the read sequence. The resulting BAM files were processed using the *jgi_summarize_bam_contig_depths* script of MetaBAT2 (v2.12.1) (50) to compute within- and between-sample coverages for each scaffold. The scaffolds were binned by running MetaBAT2 on all samples individually (--minContig 2000 and --maxEdges 500). Metagenomic bins were annotated with Anvio (v7.1.0) (51), quality-controlled using the CheckM (v1.0.13) (52) lineage workflow (completeness ≥ 50% and contamination < 10%) to generate 505 MAGs. Complete genes were predicted using Prokka (v1.14.6) (53).

Genes were subsequently clustered at 95% identity, keeping the longest sequence as representative using CD-HIT (v4.8.1) with the parameters -c 0.95 -M 0 -G 0 -aS 0.9 -g 1 -r 0 -d 0 - b 1000. Representative gene sequences were aligned against the KEGG database (release April.2022) using DIAMOND (v2.0.15) (54) and filtered to have a minimum query and subject coverage of 70% and requiring a bitScore of at least 50% of the maximum expected bitScore (reference against itself).

Quality-controlled metagenomic and metatranscriptomic sequencing reads were aligned against the set of representative genes. Gene-length normalized read abundances (i.e., gene copies) were first summarized into KEGG orthologous groups (55) and then normalized by the median of single-copy marker genes copies (56) to calculate mean per-cell gene and transcript copy numbers of orthologous groups, respectively.

### Statistical analyses

To test the hypotheses that dark CO_2_ fixation (I) increases with decreasing O_2_ availability and (II) increases with increasing salinity, we performed Spearman’s rank correlations to investigate changes in gene- and transcript abundance along the salinity gradient (ranking from salt via brackish to fresh) and the flooding gradient (ranking from high marsh via low marsh to pioneer zone). We used averages of the samples originating from the same soil core (i.e., average of deep and surface samples). For investigating changes along the depth gradient (two levels: surface vs. deep), we performed a paired t-test.

### Data availability

Raw reads of the metagenomic, metatranscriptomic and 16S rRNA analysis were deposited at the European nucleotide archive ENA under project accession number PRJEB54081 (Biosamples: SAMEA110291994 - SAMEA110292011 and SAMEA113954754 - SAMEA113954807). The gene catalog for the metagenomic and metatranscriptomic analyses and ASV tables for the 16S rRNA analyses were deposited at the ZFDM repository of the University of Hamburg (https://doi.org/10.25592/uhhfdm.13933).

## ACKNOWLEDGEMENTS

This study was funded by the Deutsche Forschungsgemeinschaft (DFG, German Research Foundation) within the Research Training Group2530: “Biota-mediated effects on Carbon cycling in Estuaries” (project number 407270017; contribution to Universität Hamburg and Leibniz-Institut für Gewässerökologie und Binnenfischerei im Forschungsverbund Berlin e.V. (IGB)).

This work was supported by the DFG Research Infrastructure NGS_CC (project 407495230) as part of the Next Generation Sequencing Competence Network (project 423957469) and core funding from ETH Zürich to the Microbiome Research Laboratory headed by S.S. Meta-transcriptome and 16S rRNA sequencing were carried out at the Competence Centre for Genomic Analysis (Kiel).

Peter Mueller was supported by the Deutsche Forschungsgemeinschaft (DFG; German Research Foundation) in the framework of the Emmy Noether Program (502681570).

## AUTHOR CONTRIBUTION STATEMENT

L.G. and W.R.S. designed the study and wrote the manuscript. H.J.R. led the metagenomic and metatranscriptomic data processing. P.M. contributed to the development of the conceptual framework and developed the statistical approach. H.-P.G., P.M and J.N.W. edited the manuscript. S.S. contributed computational resources. L.G. and A.T. performed laboratory work and analyzed the data. J.N.W. contributed to the analysis of data. All authors reviewed the manuscript.

## COMPETING INTEREST

The authors declare to have no financial or non-financial conflict of interest.

---

**Figure S1:**
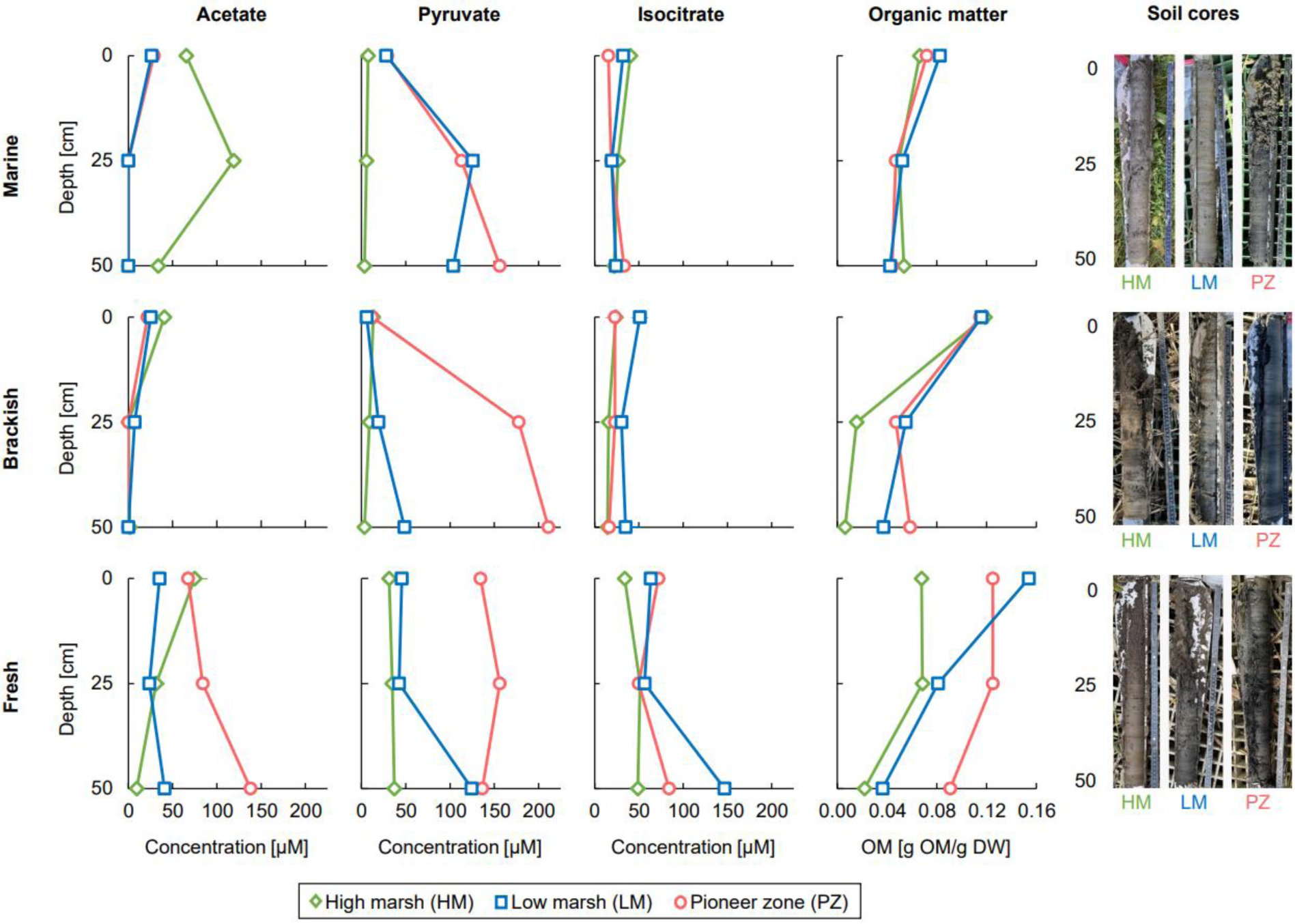
Depth profiles showing abundance of metabolites and organic matter in the salt marsh, the brackish marsh and the freshwater marsh each for the high marsh (green), the low marsh (blue) and the pioneer zone (red). The panels in the first three columns show the concentrations of acetate, pyruvate and isocitrate. The measurements were conducted in duplicates each of the same environmental sample. Error bars are too small to be visible for most of the sites. These metabolites are key components in CO_2_ fixing pathways, such as the WLP and rTCA cycle. Pyruvate plays a significant role in carbon metabolism, driving the TCA cycle and promoting oxidative phosphorylation. Acetate is formed under anaerobic conditions and can be produced by the WLP, rTCA cycle, and fermentation. Isocitrate is an intermediate in both the TCA and rTCA cycles, as well as in other pathways. Concentrations of the organic acids varied from 0 to 211 µM across all locations, without a clear pattern related to soil layers or depth. Acetate showed high concentrations, particularly in the freshwater marsh (137.8 ± 2.3 µM (mean ± SD)), while highest pyruvate concentrations were measured mainly in the low marsh and the pioneer zones of all three sites along the salinity gradient (211.1 ± 1.1 µM). Isocitrate exhibited the highest concentrations in the freshwater marsh (34.0 ± 8.0 µM to 146.6 ± 1.0 µM). The fourth column shows the amount of the soil organic matter, which serves as a crucial variable for microbial function and biogeochemistry in wetlands. To assess its impact, we quantified organic matter content as a proxy for substrate availability in our microbial and metabolic analyses. Organic matter contents ranged from 0.01 to 0.15 g OM/g DW. Salt marsh samples showed relatively constant organic matter amounts (0.04-0.08 g OM/g DW) across flooding and soil depth. Brackish marsh samples exhibited greater variation (0.01-0.12 g OM/g DW), while freshwater marsh samples ranged from 0.02 to 0.15 g OM/g DW. Surface values were 2-19 times higher than values of the deeper layers, confirming the organic matter availability gradient. The last column shows pictures of the respective soil cores used for the metabolite and organic matter analyses.

[added as supplementary Excel file]

**Table S1:** Bioinformatic output summary of metagenome-(METAG) and metatranscriptome (METAT) samples.

[added as supplementary Excel file]

**Table S2:** List of all observed phyla and classes with respective counts in 16S amplicon dataset.

[added as supplementary Excel file]

**Table S3:** Taxonomic assignment to metagenome-(METAG) and metatranscriptome (METAT) reads for the nine CO_2_ fixation related key genes.

[added as supplementary Excel file]

**Table S4:** Assignment of the most abundant microbial groups to the nine genes associated with dark CO_2_ fixation based on MAGs. Numbers indicate percentage of the genes (METAG) / transcripts (METAT) assigned to the respective microbial group at the 18 different sites. Last four columns give the average abundance [%] for all sites with a gene/transcript abundance >0 and the standard deviation.

## Notes

The authors declare no conflict of interest.

https://doi.org/10.25592/uhhfdm.13933

https://www.ebi.ac.uk/ena/browser/view/PRJEB54081

### Competing Interest Statement

The authors have declared no competing interest.

